# Respiration enhances TDP-43 toxicity, but TDP-43 retains some toxicity in the absence of respiration

**DOI:** 10.1101/415893

**Authors:** Sei-Kyoung Park, Sangeun Park, Susan W. Liebman

**Affiliations:** Department of Pharmacology, University of Nevada, Reno NV, USA

**Author notes:** Corresponding author: Susan W. Liebman, 1664 N. Virginia St. Mail Stop 318, Reno, 89557.

## Abstract

The trans-activating response DNA-binding protein 43 (TDP-43) is a transcriptional repressor and splicing factor. TDP-43 is normally mostly in the nucleus, although it shuttles to the cytoplasm. Mutations in TDP-43 are one cause of familial amyotrophic lateral sclerosis (ALS). In neurons of these patients, TDP-43 forms cytoplasmic aggregates. In addition, wild-type TDP-43 is also frequently found in neuronal cytoplasmic aggregates in patients with neurodegenerative diseases not caused by TDP-43 mutations. TDP-43 expressed in yeast causes toxicity and forms cytoplasmic aggregates. This disease model has been validated because genetic modifiers of TDP-43 toxicity in yeast have led to the discovery that their conserved genes in humans are ALS genetic risk factors. While how TDP-43 is associated with toxicity is unknown, several studies find that TDP-43 alters mitochondrial function. We now report that TDP-43 is much more toxic when yeast are respiring than when grown on a carbon source where respiration is inhibited. However, respiration is not the unique target of TDP-43 toxicity because we found that TDP-43 retains some toxicity even in the absence of respiration. We found that H_2_O_2_ increases the toxicity of TDP-43, suggesting that the reactive oxygen species associated with respiration could likewise enhance the toxicity of TDP-43. In this case, the TDP-43 toxicity targets in the presence or absence of respiration could be identical, with the reactive oxygen species produced by respiration activating TDP-43 to become more toxic or making TDP-43 targets more vulnerable.

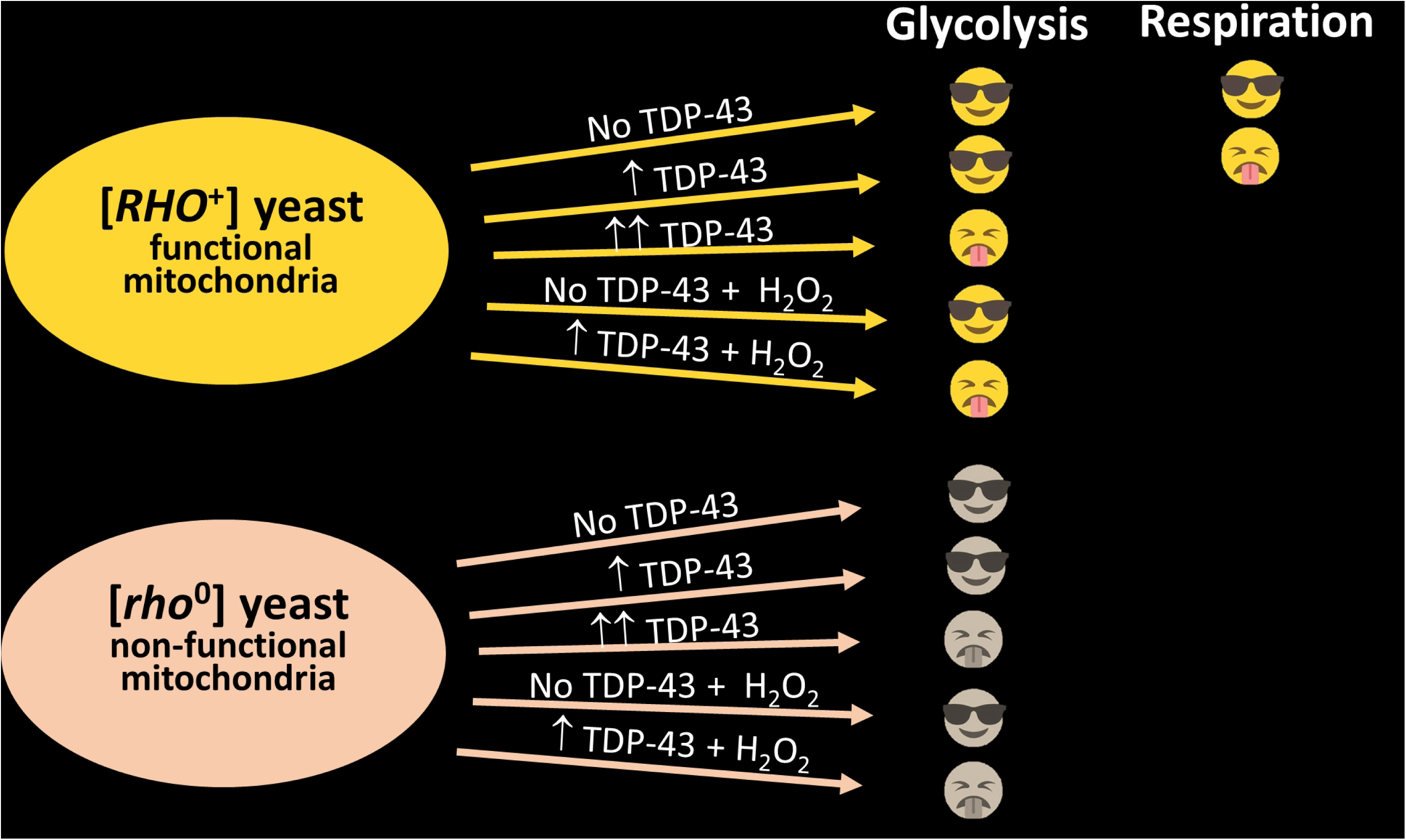

## Communication

The trans-activating response DNA-binding protein 43 (TDP-43) is a nucleic acid binding protein that functions as a transcriptional repressor, splicing factor and in translational regulation. TDP-43 is normally found mostly in the nucleus, although it shuttles between the nucleus and the cytoplasm. Mutations in *TARDBP*, the gene encoding TDP-43, are one cause of familial amyotrophic lateral sclerosis (ALS) and frontotemporal dementia (FTD). In neurons of these patients, TDP-43 is no longer found in the nucleus, but instead forms cytoplasmic aggregates. In addition, wild-type TDP-43 is also frequently found in cytoplasmic aggregates in neurons of patients with other neurodegenerative diseases or with ALS/FTD not caused by TDP-43 mutations^1; 2; 3; 4^.

How TDP-43 aggregates are associated with toxicity is the subject of intense research. Evidence links TDP-43 toxicity with both inhibition of the ubiquitin proteasome system^5^, and inhibition of lysosome and endosomal activity^6; 7^. Also, several studies find that TDP-43 alters mitochondrial function ^8; 9; 10; 11; 12; 13; 14; 15^. Overexpression of TDP-43 in a number of model organisms causes neurodegeneration similar to that seen in ALS patients. In motor neurons of TDP-43 transgenic mice, mitochondria were found in cytoplasmic inclusions or in abnormal juxta-nuclear aggregates and were missing in motor axon termini^9; 10; 11^. In mammalian neuron-like cell culture, TDP-43 localized to mitochondria and caused mitophagy^12^. Likewise, in a mouse model, TDP-43 co-localization with motor neuron mitochondria was enhanced by TDP-43 ALS mutations and this co-localization was associated with inhibited mitochondrial function^13; 14^. Also, overexpressed TDP-43 in human cell culture binds to, and inhibits maturation of, mitochondrial RNA causing a phenotype similar to that seen in cells deficient in mitochondrial RNase P^15^. Finally, mutations in the mitochondrial intermembrane protein, CHCHD10 (yeast homolog MIX17) have recently been associated with sporadic and familial ALS and FDT ^16; 17; 18; 19; 20; 21; 22^ and functional CHCHD10 appears to help keep TDP-43 in the nucleus away from mitochondria^23^.

There is also evidence that TDP-43 expression in yeast causes mitochondrion-dependent apoptosis and that TDP-43 toxicity requires respiratory capacity^8^. TDP-43 expressed in yeast causes toxicity and forms cytoplasmic aggregates^24^. Furthermore, conserved genetic modifiers of this toxicity have been validated in higher organisms and have led to the discovery of human ALS genetic risk factors^25^. This confirms the usefulness of the yeast model.

To examine the effect of respiration on TDP-43 toxicity we used the fact that yeast turns off respiration in dextrose media where it instead uses glycolysis to grow^26^. We found that TDP-43-GFP expressed with a *TET* promoter aggregated in yeast cells whether grown on dextrose, where respiration is inhibited; galactose, where respiration is not inhibited and both fermentation and respiration is present ^26^; or glycerol or ethanol where only respiration and not fermentation is present. However, the *TET* controlled TDP-43 was only toxic on galactose, glycerol or ethanol where respiration is present. Also, the previously described TDP-43 aggregation and induction of cell elongation^5^ was much more pronounced on media with respiration (galactose and glycerol) vs. media without respiration (dextrose) (Fig. 1ab). Despite this, there was no significant difference in the level of soluble TDP-43 found in boiled lysates of dextrose vs. galactose or glycerol grown cells (Fig. 1c). To test if the difference in sensitivity to TDP-43 was caused by the slower cell division rate in galactose vs. dextrose, we compared the toxicity in cells grown on dextrose at room temperature with cells grown on galactose at 30°C. Although cells grew at approximately the same rate under these two conditions, TDP-43 toxicity was pronounced on galactose but not dextrose plates (Supplementary Fig. 1).

**Fig. 1.**
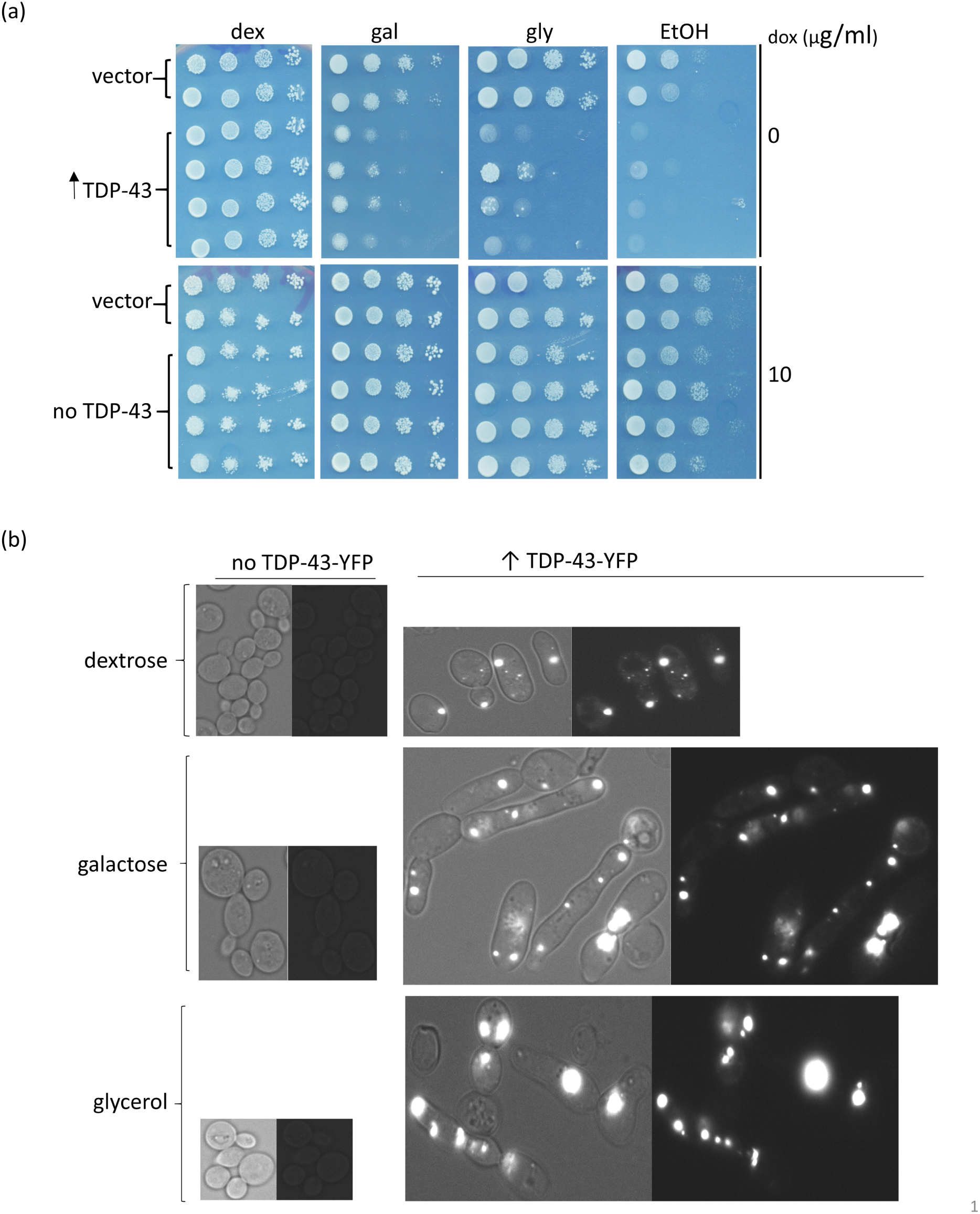
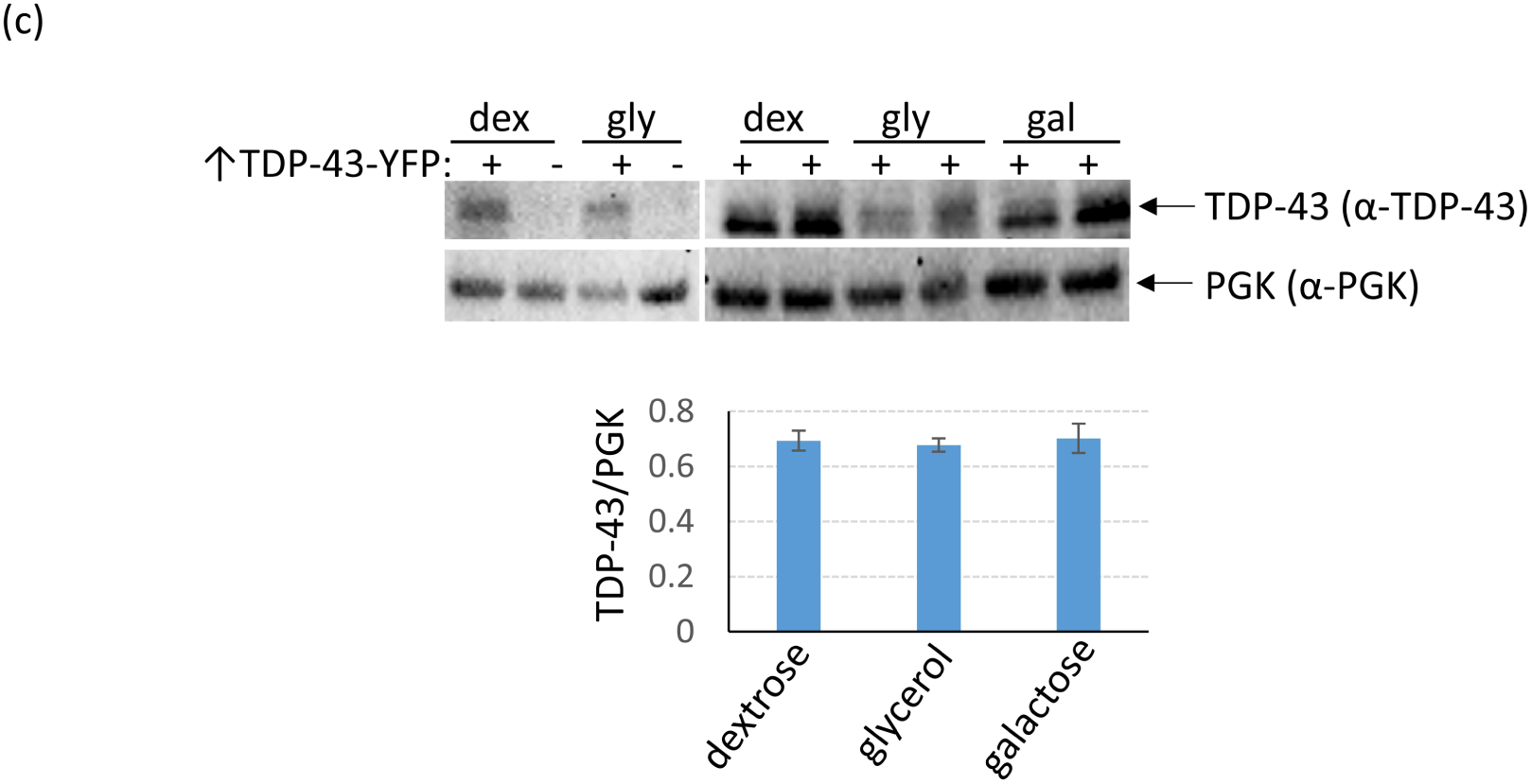
TDP-43 overexpression causes more toxicity and cell elongation in cells grown in media that allows respiration vs. cells grown in the absence of respiration. (a) Yeast cells (L2910 [*pin*^*-*^] [*RHO*^*+*^] 74D-694) transformed with the doxycycline-repressible *CEN TET-TDP-43-YFP TRP1* (p2223), or vector control (p1576) plasmid were obtained on plasmid selective dextrose plates with 10 µg/ml of doxycycline (dox) to inhibit expression of TDP-43-YFP. Normalized suspensions of cells were serially diluted in water (1/10) and were spotted on plasmid selective 2% dextrose (dex), 2% galactose (gal), 2% glycerol (gly), and 2% ethanol (EtOH) plates without doxycycline (0) and with 10 µg/ml of doxycycline to respectively express or not express TDP-43-YFP. The plates were photographed after 4 (dextrose) or 6 (galactose, glycerol or ethanol) d of incubation at 30°C. (b) TDP-43 aggregates in galactose, glycerol and dextrose but causes more aggregation and cell elongation in galactose or glycerol vs. dextrose media. Cells from TDP-43-YFP inducing plates (↑TDP-43-YFP) were imaged with a fluorescence microscope at the same exposure and are shown at the same magnification. Left images show bright field with fluorescent, right images show just fluorescence. Control cells with no TDP-43-YFP expression (no TDP-43-YFP) are also shown. The increased aggregation and elongation on galactose and glycerol caused by TDP-43-YFP expression was very obvious. (c) Expression of *TET* controlled TDP-43-YFP is not enhanced on glycerol or galactose medium. The level of TDP-43-YFP was compared in L2910 transformants with doxycycline-repressible -*TDP-43-YFP* (p2223) grown in 2% dextrose (dex) vs. 2% glycerol (gly) or 2% raffinose + 2% galactose (gal) plasmid selective media. Cells grown overnight in medium containing 8 µg/ml of doxycycline were diluted to OD_600_ = 0.5 in plasmid selective dex, gly or gal without doxycycline, grown for 24 h, harvested, and lysed for immunoblotting^35^. Normalized cell lysates boiled for 5 mins with 2% SDS in 80 mM DTT sample buffer were resolved on 10% SDS-PAGE followed by immunoblotting with rabbit polyclonal a-TDP-43 antibody (1:3000, Proteintech Group). Also, PGK, yeast 3-Phosphoglycerate Kinase, was detected with anti PGK antibodies (1:10,000, Novex) as an internal control (3c upper). Immunoblot signals for TDP-43-YFP and PGK were quantified and converted into % ratios of TDP43-YFP to PGK (3c lower). Standard error shown was calculated from 3 independent immunoblots. Gateway destination vector p1576 was made by dropping reading frame B gateway cassette (http://www.lifetechnologies.com) into pCM184^36^. TDP-43-YFP from entry vector pDONR TDP-43-YFP (Addgene plasmid # 27470) was then cloned into p1576 using an LR reaction, creating doxycycline-repressible *CEN TET-TDP-43-YFP TRP1* (p2223).

One explanation could be that TDP-43 only inhibits respiration, which only affects growth when cells are respiring. Alternatively, respiration could be required for TDP-43 to be toxic, e.g. by modifying the TDP-43 protein. Indeed, oxidative stress has been shown to increase disulfide cross-linking, acetylation, and aggregation of TDP-43 in mammalian cells^27^. Furthermore, oxidative stress has been proposed to promote acetylation of TDP-43, causing reduced binding of TDP-43 to RNA and increased levels of cytoplasmic, phosphorylated, aggregated TDP-43^28^.

However, we eliminated both of the above hypotheses because we were able to detect TDP-43 toxicity in the absence of respiration on dextrose by expressing TDP-43-YFP with the strong *CUP1* promoter ^29; 30^. To do this, we obtained an array of integrants of a plasmid expressing *CUP1*-TDP43-YFP in [*PIN*^+^] 74D-694. As expected, due to differences in the number of tandem plasmid repeats present ^31^, different integrants showed different levels of toxicity (high, medium and low). The more toxic integrants clearly showed toxicity on dextrose (Supplementary Fig. 2). Each integrant was cured of [*PIN*^+^] by growth on guanidine HCl ^32^. Then, since yeast lacking functional mitochondria are viable we made all the [*PIN*^+^] and [*pin*^−^] [*RHO*^+^] integrants containing functional mitochondria into [*rho*^*0*^] cells lacking functional mitochondrial. We then compared the level of toxicity caused by overexpressed TDP-43 in these isogenic [*RHO*^+^] and [*rho*^*0*^] strains grown on dextrose (dex) where respiration is inhibited. We found that TDP-43 was toxic and retained the same level of expression in both [*RHO*^+^] and [*rho*^0^] cells (Fig. 2abc). There was also no obvious difference in the appearance of TDP-43 aggregates (Fig. 2d). As we reported previously^5^ TDP-43 is more toxic in the presence of the [*PIN*^+^] prion. This was clear when cells were grown on dextrose or galactose (Fig. 2a), but the effect of [*PIN*^+^] on TDP-43 toxicity in cells grown on glycerol was minor and not reproducible (not shown). Since the [*rho*^0^] cells can never respire and the [*RHO*^+^] cells do not respire on dextrose, this clearly establishes that there is a TDP-43 toxicity target in yeast distinct from respiration and that respiration is not required for this TDP-43 toxicity.

**Fig. 2.**
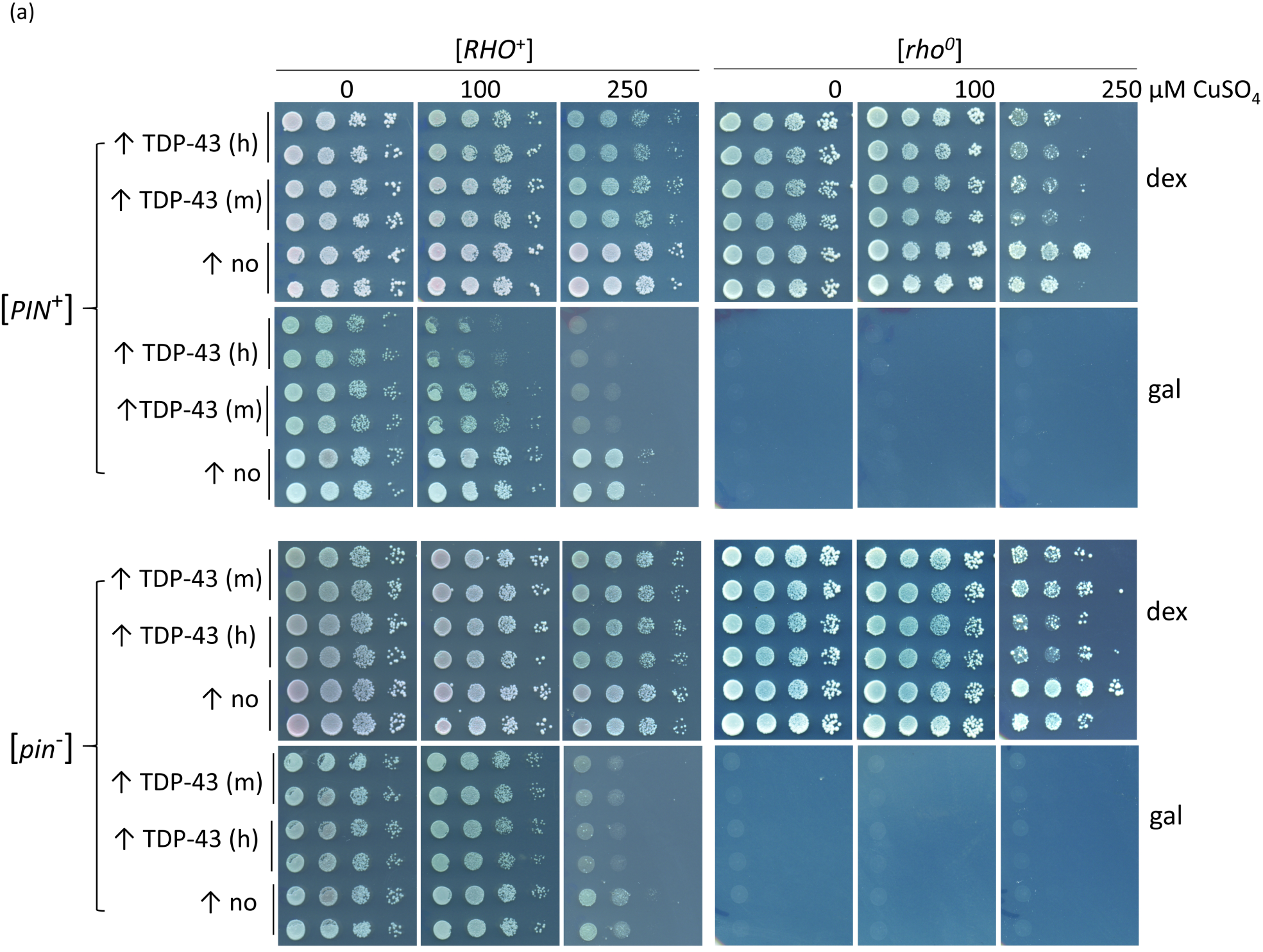
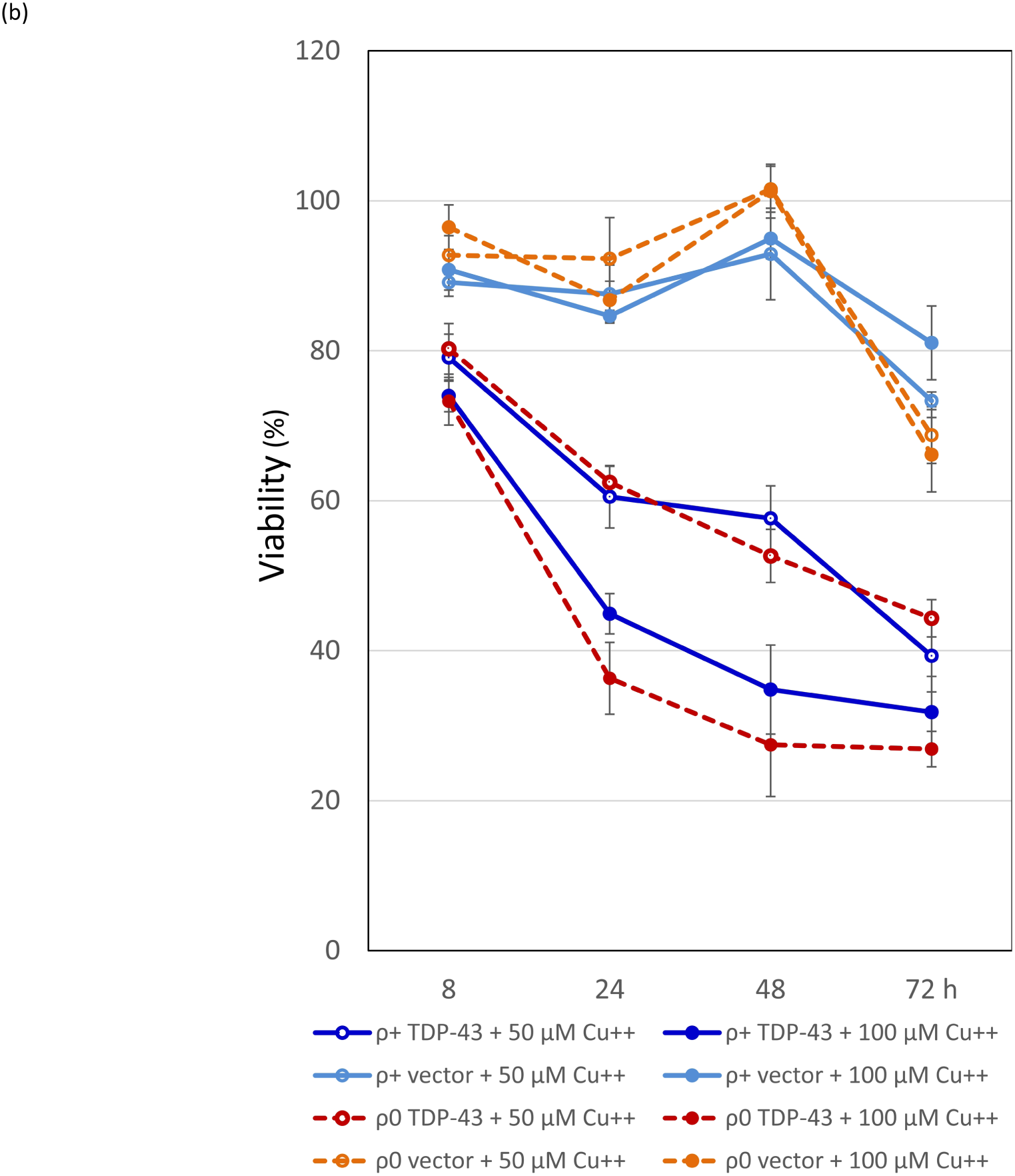
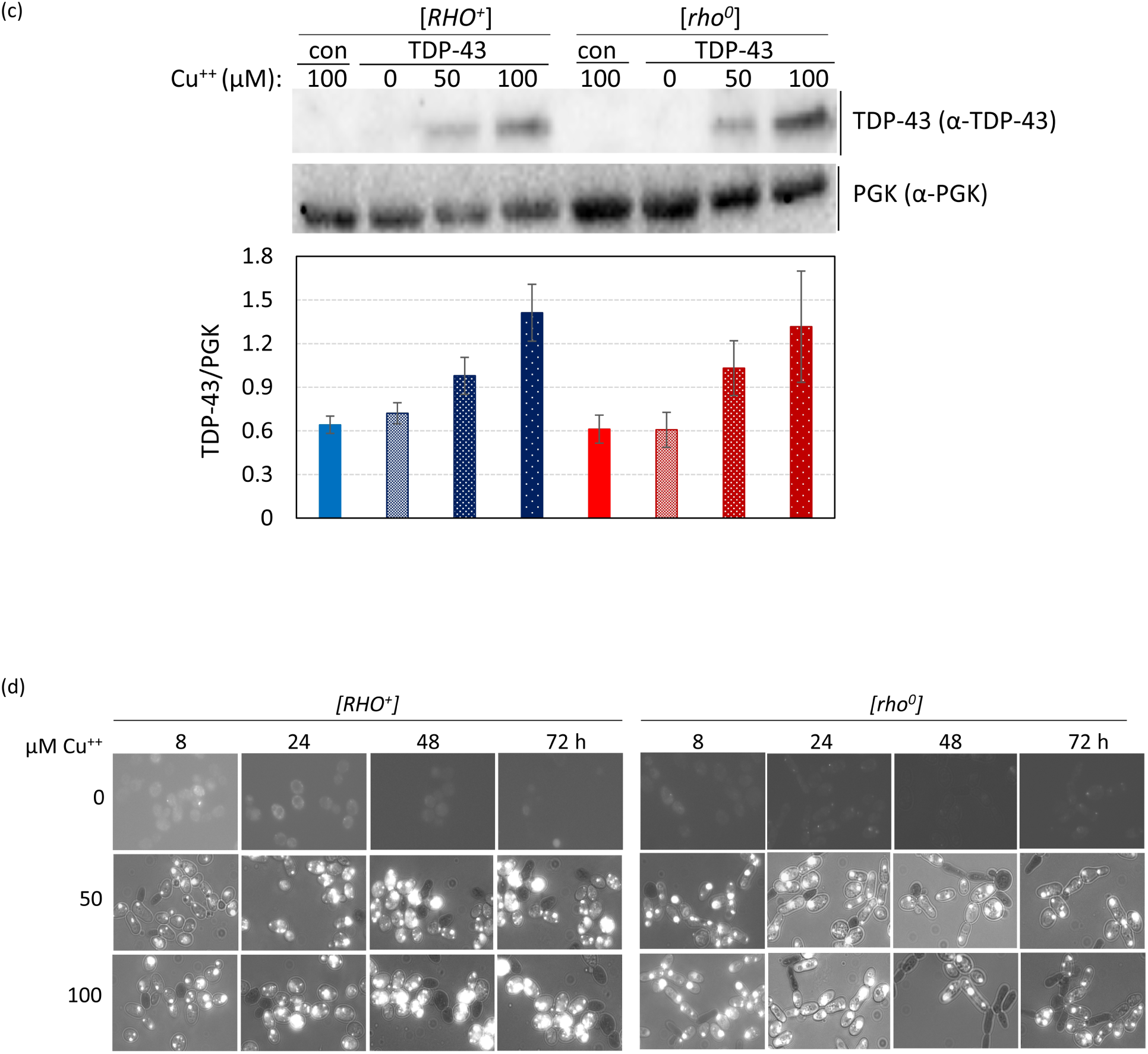
TDP-43-YFP is still toxic in non-respiring cells and in cells lacking functional mitochondria. (a) Spot test measure of TDP-43-YFP toxicity. *CUP1-TDP-43-YFP* in p2261 (pAG305 *CUP1::TDP43-YFP*) was integrated at the *LEU2* locus in [*RHO*^+^] [*PIN*^+^] and [*RHO*^+^] [*pin*^−^] 74D-694 cells. Isogenic [*rho*^*0*^] versions of the *CUP1-TDP-43-YFP* integrants were acquired by growing [*RHO*^+^] integrants on YPD with 0.4 mg/ml ethidium bromide. This method has been shown to cause the loss of all mitochondrial DNA^37^. The lack of functional mitochondria was verified as cells failed to grow on glycerol medium. To assay for TDP-43-YFP toxicity, two independent subclones of [*RHO*^+^] and [*rho*^*0*^] *CUP1-TDP-43-YFP* medium (m) and high (h) level toxicity integrants (Supplementary Fig. 2) (↑TDP-43) and control empty integrants (↑no) grown on dextrose medium without CuSO_4_ were normalized, serially diluted (1/10) and spotted on dextrose (dex) and galactose (gal) plates including 0, 100, and 250 µM CuSO_4_. Plates with [*rho*^0^] cells were allowed to grow longer than plates with [*RHO*^+^] cells and galactose plates were grown longer than dextrose plates prior to scanning. Reduced growth indicated toxicity. As expected [*rho*^*0*^] cells failed to grow on gal. Integrants were constructed by transforming cells with purified linear integrating vector p*CUP1-TDP-43-YFP* (p2261) that had been cut in the *LEU2* gene with BstXI and control integration vector without TDP-43-YFP, pAG305 GAL1-ccdB (Addgene plasmid #14137) that was cut similarly, and selecting for transformants on integrant selective dextrose plates. Integration was verified by observing TDP-43-YFP aggregation in transformants in dextrose medium with 50 µM CuSO_4_. To generate p2261, the HpaI-PmeI fragment containing the *TET* promoter from pAG305 *TET-TDP-43-YFP* (p2237) was switched with the *CUP1* promoter from p1988 (*pCUP1-SOD1-GFP*) cut with the same restriction site. Plasmid pAG305 *TET-TDP-43-YFP* (p2237) was made by replacing the *GAL* promoter in pAG305 *GAL-ccdB* (Addgene plasmid # 14137) with PCR amplified HpaI-XbaI *TET* promotor from pCM184. Integrants contain different numbers of tandem plasmids which caused different levels of toxicity. (b) Viability measure of TDP-43-YFP toxicity. Cytotoxicity of the *CUP1* driven TDP-43-YFP in both [*RHO*^+^] and [*rho*^*0*^] cells was determined by colony forming units^8^. Overnight cultures of a TDP-43-YFP integrant associated with a high level of toxicity (supplementary Fig. 2) or vector controls grown without CuSO_4_ were inoculated into liquid synthetic dextrose media with 0, 50 or 100 µM CuSO_4_ to a final OD_600_ of 0.5. Samples were taken thereafter at 8, 24, 48, and 72 h and colony forming units determined on synthetic dextrose plates lacking CuSO_4_. The number of colonies grown after 3 d of incubation at 30°C were counted and converted into viability (%) calculated as ratio of cells grown in TDP-43-YFP inducing media (with 50 or 100 µM CuSO_4_) over cells grown in non-inducing media (0 µM CuSO_4_). Error bars present the standard error calculated from 3 independent subclones of integrants. (c) No significant difference in levels of TDP43-YFP expression in [*RHO*^+^] vs. [*rho*^*0*^] *CUP1-TDP-43-YFP* integrated strains. Isogenic [*RHO*^+^] vs. [*rho*^*0*^] high toxicity integrants taken from dextrose plates were inoculated into 50 ml dextrose containing 0, 50, or 100 µM CuSO_4_ to an OD_600_ of 0.5. Cells were harvested and lysed at 24 h. Normalized cell lysates were boiled, resolved and immunoblotted as shown in Fig. 1c. The intensity of each protein band (upper) was scanned and the ratio between TDP-43-YFP and PGK was calculated (lower). Bars represent standard error calculated from 3 independent immunoblots. (d) TDP-43-YFP aggregates similarly in [*RHO*^+^] and [*rho*^0^] *CUP1-TDP-43-YFP* integrated strains. Samples from 2b were visualized with fluorescent microscope at the same exposure. No obvious visual differences were seen.

To test the idea that increased levels of reactive oxygen species associated with respiration could be the cause of the observed increase in TDP-43 toxicity in the presence of respiration, we looked at the effects of H_2_O_2_, a source of reactive oxygen species, and N-Acetyl L-cysteine (NAC) an antioxidant, on TDP-43 toxicity. Using an integrant with low TDP-43 toxicity, we saw no toxicity of cells grown for 24 h with either 2mM H_2_O_2_ without TDP-43 expression (Fig. 3a left) or with expression of TDP-43 with 250 μM CuSO_4_ in dextrose medium without H_2_O_2_ (Fig. 3a right rows marked 0). However, the combination of 24 h of 2mM H_2_O_2_ and induction of TDP-43 with 250 μM CuSO_4_ did cause toxicity (Fig. 3a right row marked 2). This was true in both [*RHO*^+^] or [*rho*^0^] cells. We also observed a dramatic increase in the size of TDP-43 aggregates in cells treated with H_2_O_2_ (Fig. 3b), and this was not due to an increase in the cellular level of TDP-43 (Fig. 3c). Possibly, this was caused by free radical oxygen stress, which has been shown to cause an increase in protein aggregation^33; 34^. Also, we saw reduced toxicity of cells in the presence of the antioxidant NAC and this was quite dramatic in the presence of TDP-43 (Fig. 4ab). However, the level of TDP-43 in cells was dramatically reduced in the presence of NAC (Figs. 3c and 4c). If NAC directly reduced expression of TDP-43 that would explain the reduced toxicity. Likewise if NAC inhibited aggregation of TDP-43, the unaggregated TDP-43 would be expected to be rapidly degraded again explaining reduced toxicity.

**Fig. 3.**
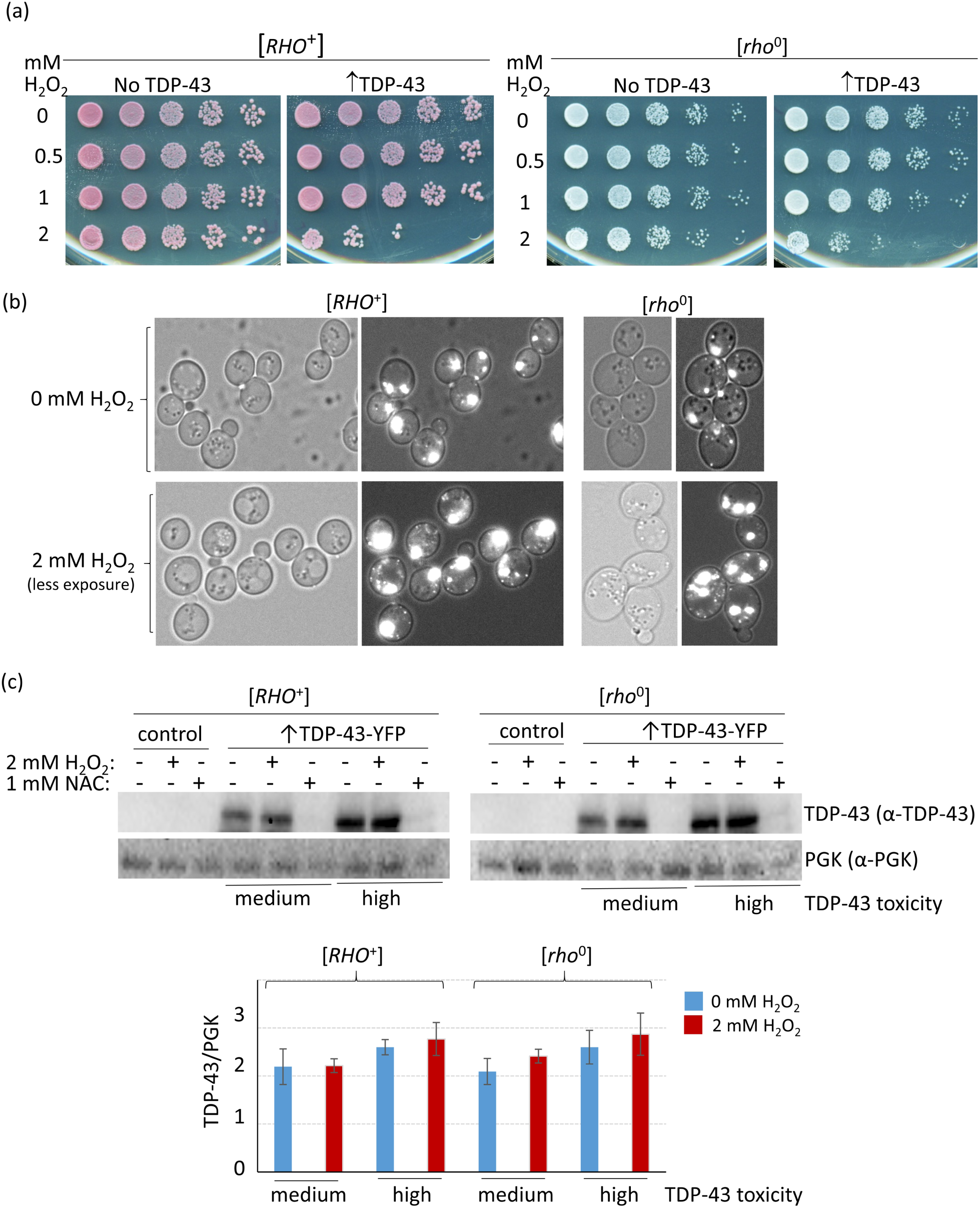
H_2_O_2_ enhances TDP-43 toxicity and aggregation. (a) H_2_O_2_ enhances TDP-43 toxicity. [*RHO*^*+*^] *and* [*rho*^*0*^] versions of strain 74D-694 with integrated *CUP1-TDP-43-YFP* (↑TDP-43) associated with a low level of toxicity (Supplementary Fig. 2), or vector control (No TDP-43), were grown to OD_600_ = 0.5 in synthetic dextrose non-inducing medium and then for an additional 24 h after the simultaneous addition of CuSO_4_ to 250 μM to induce TDP-43-YFP expression and H_2_O_2_ to the concentrations listed. Serial dilutions of cells were then spotted on plates lacking CuSO_4_ and H_2_O_2_ and photographed after growth. (b) H_2_O_2_ enhances TDP-43 aggregation. Cells from the liquid cultures used in (a) were examined under a fluorescent microscope. Left shows bright field, right is fluorescence plus bright field. Since the intensity of TDP-43-YFP aggregates was much brighter in H_2_O_2_ treated cells than in cells without H_2_O_2_ treatment, images of H_2_O_2_ treated cells were taken with less exposure than cells without H_2_O_2_ treatment to allow visualization of dots. (c) The level of TDP-43 in cells is unchanged by the addition of H_2_O_2_, but is dramatically reduced by the addition of NAC. Freshly grown subclones of medium and high level toxicity *CUP-TDP-43-YFP* integrants or control (empty vector) were inoculated at OD_600_ = 0.5 into 50 mL plasmid selective dextrose media with 250 µM CuSO_4_ and either 2 mM H_2_O_2_ or 1 mM NAC. Cells were harvested and lysed after 24 h growth. Normalized cell lysates were boiled, resolved and immunoblotted as shown in Fig. 1c. The protein level was compared between cells treated or not treated with H_2_O_2_ or NAC. The ratio between scanned TDP-43-YFP and PGK signals for control (blue) and H_2_O_2_ (red) cells is shown. Bars represent standard error calculated from 3 independent immunoblots.

**Fig. 4.**
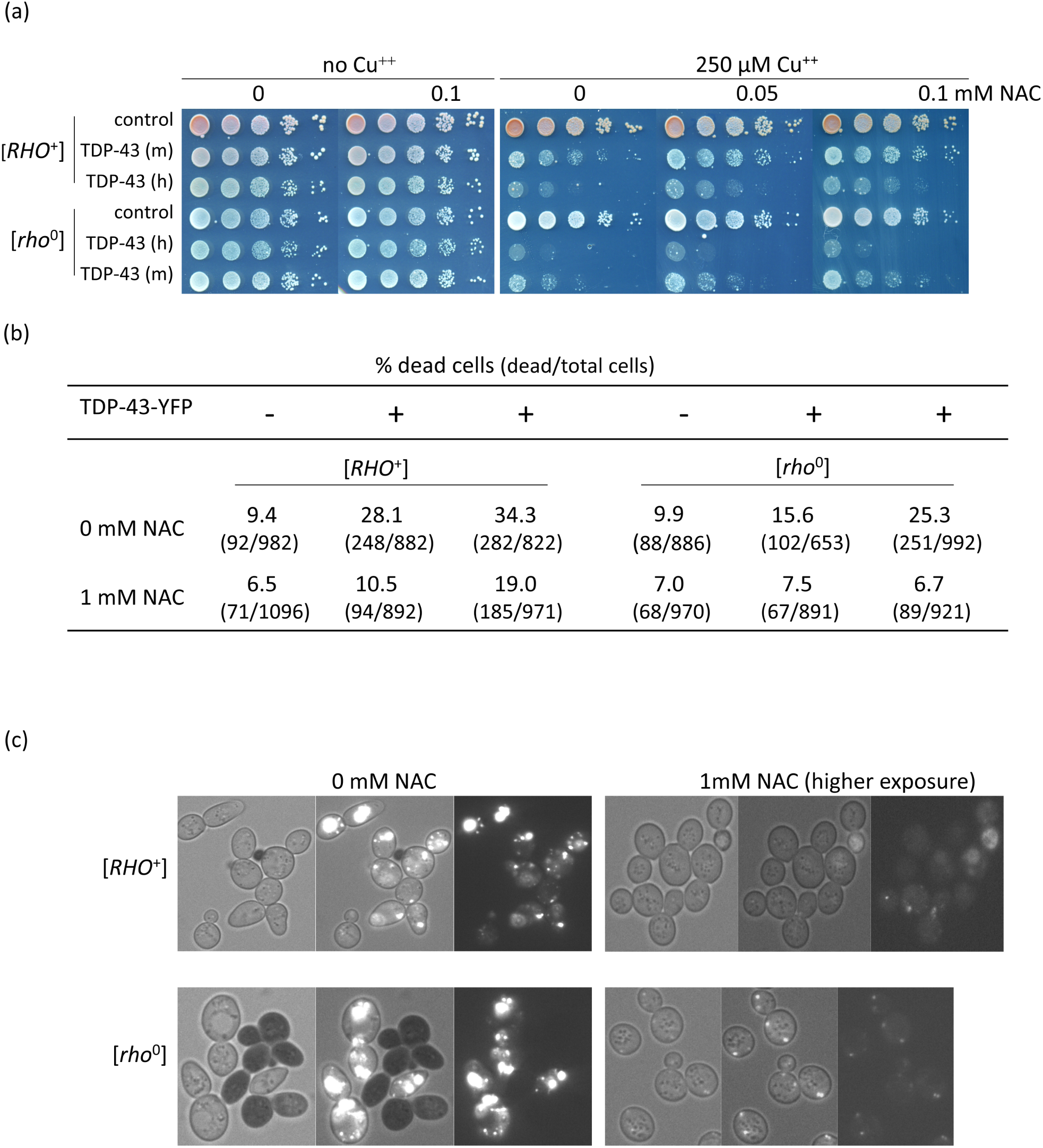
Antioxidant reduces TDP-43 level and toxicity. (a) Antioxidant reduces TDP-43 growth inhibition. Isogenic [*RHO*^*+*^] and [*rho*^*-*^] versions of *CUP1-TDP-43-YFP* (TDP-43) integrants with medium (m) and high (h) levels of toxicity and vector (control) grown on plasmid selective medium were normalized, serially diluted (1/10) and spotted on plates with the indicated levels of CuSO_4_ and antioxidant, NAC (N-acetyl-L-cysteine, Sigma). Plates were photographed after 4 days of incubation at 30°C. (b) NAC reduces lethality due to TDP-43. An integrant of *CUP1-TDP-43-YFP* (TDP-43) with high toxicity and vector (control) grown on synthetic dextrose medium without CuSO_4_ were inoculated to OD_600_ = 0.5 in dextrose medium with (+) or without (−) 250 µM CuSO_4_ and 0 mM or 1 mM NAC. After 24 h growth cells were stained with trypan blue to identify dead cells^5^ and counted. (c) NAC reduces TDP-43-YFP level. Cells from (b) were photographed with bright field (left), fluorescence and bright field (middle) and just fluorescence (right). TDP-43-YFP fluorescence was barely observed in NAC treated cells even at the very high exposure shown. Black cells indicate trypan blue stained dead cells.

The prevalence of TDP-43 aggregation in patients with a variety of neurodegenerative diseases makes it critical to understand how this is associated with toxicity. Our data shows that toxicity is enhanced in the presence of respiration, but that TDP-43 remains toxic even in the absence of respiration. This is consistent with the hypothesis that TDP-43 targets the same cellular components in the presence or absence of respiration, and that the reactive oxygen species produced by respiration either activates TDP-43 to become more toxic or makes the same TDP-43 targets more vulnerable.

## Acknowledgements

We thank Aaron Gitler, Stanford U. for plasmids and Ruben Dagda, U. of Nevada, Reno, and Martin Duennwald, Western U. Ca, for helpful ideas. This work was supported National Institutes of Health Grant R01GM056350 (SWL).

---

**Supplementary Fig. 1.**
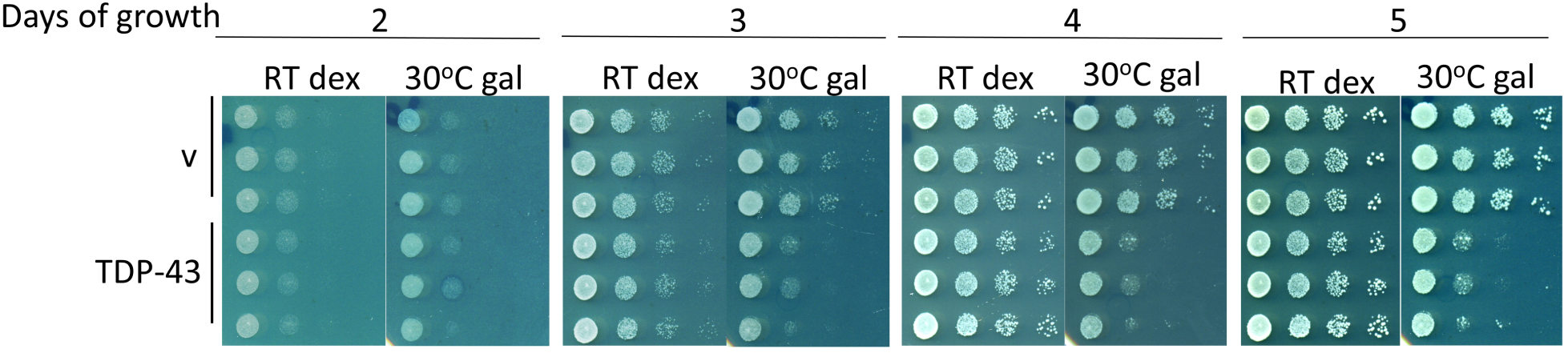
Slow growth does not enhance TDP-43 toxicity. Three independent transformants of [*PIN*^*+*^] 74D-694 (L1749) with either *TET-TDP-43-YFP* (TDP-43) or empty vector (v) were taken from plasmid selective media without TDP-43-YFP expression, normalized to OD_600_ = 3 and serially diluted (1/10) and spotted on dextrose (dex) or galactose (gal) plasmid selective inducing media (lacking doxycycline). Dextrose and galactose plates were respectively incubated at room temperature (RT) or 30°C for the number of days indicated prior to being photographed.

**Supplementary Fig. 2.**
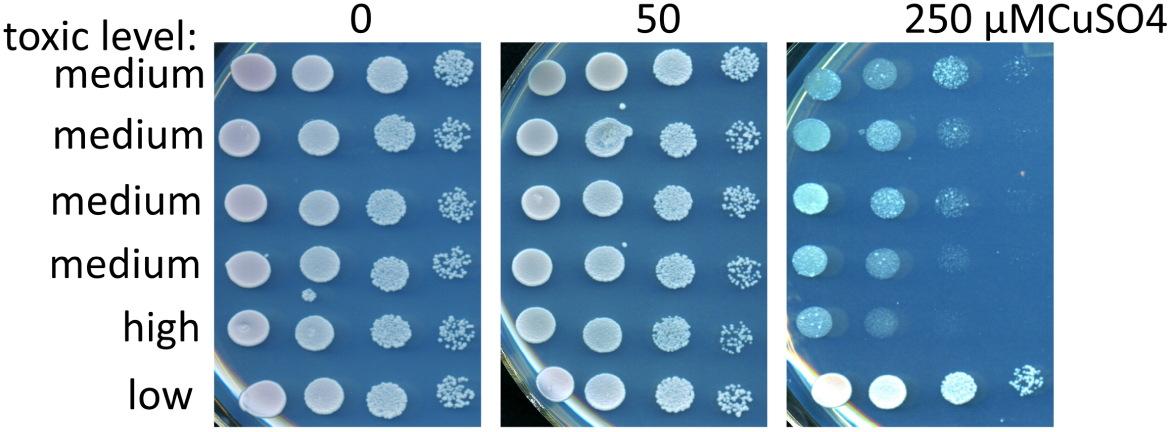
*CUP1* controlled *TDP-43-YFP* integrants show differences in toxicity. Cells of different [*PIN*^+^] integrants were normalized and spotted on plasmid selective dextrose plates containing 0, 50, and 250 µM CuSO_4_ as shown. Integrants were scored as having low, medium and high toxicity.

## Abbreviations

TDP-43: trans-activating response DNA-binding protein 43
ALS: amyotrophic lateral sclerosis
dex: dextrose
gal: galactose
dox: doxycycline
CuSO_4_: copper sulfate
H_2_O_2_: hydrogen peroxide
NAC: N-Acetyl L-cysteine

